# *Anopheles* Mosquitoes May Drive Invasion and Transmission of Mayaro Virus across Geographically Diverse Regions

**DOI:** 10.1101/359349

**Authors:** Marco Brustolin, Sujit Pujhari, Cory A. Henderson, Jason L. Rasgon

## Abstract

The Togavirus (Alphavirus) Mayaro virus (MAYV) was initially described in 1954 from Mayaro County (Trinidad) and has been responsible for outbreaks in South America and the Caribbean. Imported MAYV cases are on the rise, leading to invasion concerns similar to Chikungunya and Zika viruses. Little is known about the range of mosquito species that are competent MAYV vectors. We tested vector competence of 2 MAYV genotypes for six mosquito species (*Aedes aegypti*, *Anopheles gambiae*, *An. stephensi*, *An. quadrimaculatus*, *An. freeborni*, *Culex quinquefasciatus*). *Ae. aegypti* and *Cx. quinquefasciatus* were poor MAYV vectors, and either were poorly infected or poorly transmitted. In contrast, all *Anopheles* species were able to transmit MAYV, and 3 of the 4 species transmitted both genotypes. The *Anopheles* species tested are divergent and native to widely separated geographic regions, suggesting that *Anopheles* may be important in the invasion and spread of MAYV across diverse regions of the world.

## Introduction

Mayaro virus (MAYV) is a member of the genus *Alphavirus* (family Togaviridae) which was first isolated from the blood of five febrile workers in Mayaro County, Trinidad, in 1954 (*1*). MAYV is a single-stranded positive-sense RNA virus of approximately 11.7 kb and is classified in three genotypes: D, L, and N (*2,3*). Genotype D (dispersed) includes strains isolated in several South American countries, whereas genotype L (limited) includes strains isolated only in Brazil. In 2010, a minor genotype called N (new), was isolated in Peru, but it is limited to one known sequence. Since its first isolation, MAYV has caused sporadic outbreaks and small epidemics in several countries of South and Central America (reviewed in *4*). In 2015, the case of an 8-year-old child from Haiti co-infected with MAYV and Dengue virus (DENV) suggested that MAYV may also be actively circulating in the Caribbean (*5*). Several imported cases recently reported in the Netherlands (*6*), Germany (*7*), France (*8*), and Switzerland (*9*) highlight the need to survey naive regions, such as the United States, for possible introductions of this neglected arthropod-borne virus (arbovirus).

The symptoms of Mayaro fever (MAYF) include rash, fever, myalgia, retro-orbital pain, headache, diarrhea, and arthralgia, which may persist for months or even years (*10*), and are similar to caused by others arboviruses, such DENV or Chikungunya virus (CHIKV). Due to the absence of routine differential diagnostics, reported cases of MAYV likely underestimate the real prevalence, and the circulation of the virus can pass undetected in areas with ongoing DENV or CHIKV outbreaks (*4,11*).

MAYV is thought to be principally transmitted by the bite of diurnal canopy-dwelling mosquitoes of the genus *Haemagogus* (*4*). These mosquitoes are responsible for maintaining the sylvatic cycle involving nonhuman primates and birds as primary and secondary hosts, respectively. Human infections are sporadic, likely because *Haemagogus* spp. rarely display anthropophilic behaviors, and they possess a preference for rural areas with proximity to forests (*12*). Vector competence (VC) studies demonstrated that anthropophilic and urban-adapted species, such as *Aedes aegypti* and *Ae. albopictus,* are competent vectors for MAYV in laboratory conditions (*13,14*). *Culex quinquefasciatus* mosquitoes positive for MAYV have also been identified from field collections during a DENV outbreak in Mato Groso County, Brazil (*15*); however, their capacity to transmit MAYV has not been demonstrated.

Overall, little data is available about the VC of mosquitoes for MAYV (*16–18*) and, furthermore, there have been no studies about the VC of autochthonous vector species of the United States. To address this knowledge gap, we evaluated the ability of *Anopheles stephensi* (Liston, 1901), *An. gambiae* (Giles, 1902), *An. freeborni* (Aitken, 1939), *An. quadrimaculatus* (Say, 1824), *Cx. quinquefasciatus* (Say, 1823), and *Ae. aegypti* (Linnaeus, 1762) to become infected with and transmit MAYV after feeding on a viremic blood meal. Our results demonstrate that while *Ae*. *aegypti* and *Cx. quinquefasciatus* are poor vectors for MAYV, all tested *Anopheles* species were competent laboratory vectors for MAYV, including species that they have the potential to support the transmission cycle if the virus is introduced into the United States. Additionally, the results of our study provide useful information to improve entomologic surveillance programs and prevent future outbreaks of this emerging neglected pathogen.

## Material and Methods

Six mosquito species were used in this experimental study. The *An. gambiae* (NIH strain) were originally obtained from The National Institutes of Health (Bethesda, MD, USA). *An. stephensi* (Liston strain) were provided by Johns Hopkins University (Baltimore, MD, USA). *Cx*. *quinquefasciatus* (Benzon strain) were provided by the Wadsworth Center (Slingerlands, NY, USA) and was initially derived from a colony maintained by Benzon Research (Carlisle, PA, USA). *An. quadrimaculatus* (Orlando strain, MRA-139) and *An. freeborni* (F1 strain, MRA-130) were provided by BEI Resources (Manassas, VA, USA). *Ae. aegypti* (Rockefeller strain) were provided by Johns Hopkins University.

Mosquito colonies were reared and maintained at the Millennium Sciences Complex insectary (The Pennsylvania State University, University Park, PA, USA) at 27°C ± 1°C, 12:12 h light:dark diurnal cycle at 80% relative humidity in 30×30×30-cm cages. Ground fish flakes (TetraMin, Melle, Germany) were used to feed *Anopheles* spp. and *Aedes* sp. larvae. A 1:1:1 mixture of bovine liver powder (MP Biomedicals, Solon, OH, USA), koi pellets (TetraPond Koi Vibrance; TetraPond, Prestons, Australia), and rabbit pellets (Kaytee, Chilton, WI, USA) was used for *Culex* sp. larvae. Adult mosquitoes were provided with 10% sucrose solution *ad libitum* for maintenance. For reproduction and virus infection purposes, adults were fed with expired anonymous human blood (Biological Specialty Corporation, Colmar, PA, USA).

Two strains of MAYV were used for the experimental infections: BeAr 505411 (BEI Resources, Manassas, VA, USA), a genotype L strain isolated from *Haemagogus janthinomys* mosquitoes in Para, Brazil, in March 1991, and BeAn 343102 (BEI Resources, Manassas, VA, USA), a genotype D strain originally isolated from a monkey in Para, Brazil, in May 1978. Both viruses were passed once in African green monkey kidney (Vero) cells. Virus-infected supernatant was aliquoted and stored at −70°C until used for mosquito infections. Viral stock titers were obtained by the focus forming unit (FFU) technique, as described below.

Five-to seven-day-old females that had not previously blood-fed were used in this experiment. The mosquitoes were allowed to feed on infected human blood via a glass feeder jacketed with 37°C distilled water for 1 h. Aliquots of the infectious bloodmeals were collected and titers of MAYV were determined by FFU (Table 1). After blood feeding, mosquitoes were anesthetized and fully engorged females were selected and placed in cardboard cages. Infection rate (IR), dissemination rate (DIR), transmission rate (TR), and transmission efficiency (TE) were assessed at 7 and 14 days post-infection (dpi). IR was measured as the rate of mosquitoes with infected bodies among the total number of analyzed mosquitoes. DIR was measured as the rate of mosquitoes with infected legs among the mosquitoes with positive bodies. TR was measured as the rate of mosquitoes with infectious saliva among the mosquitoes with positives legs, and TE measured as the rate of mosquitoes with infectious saliva among the total number of analyzed mosquitoes (*19*).

**Table.1.**
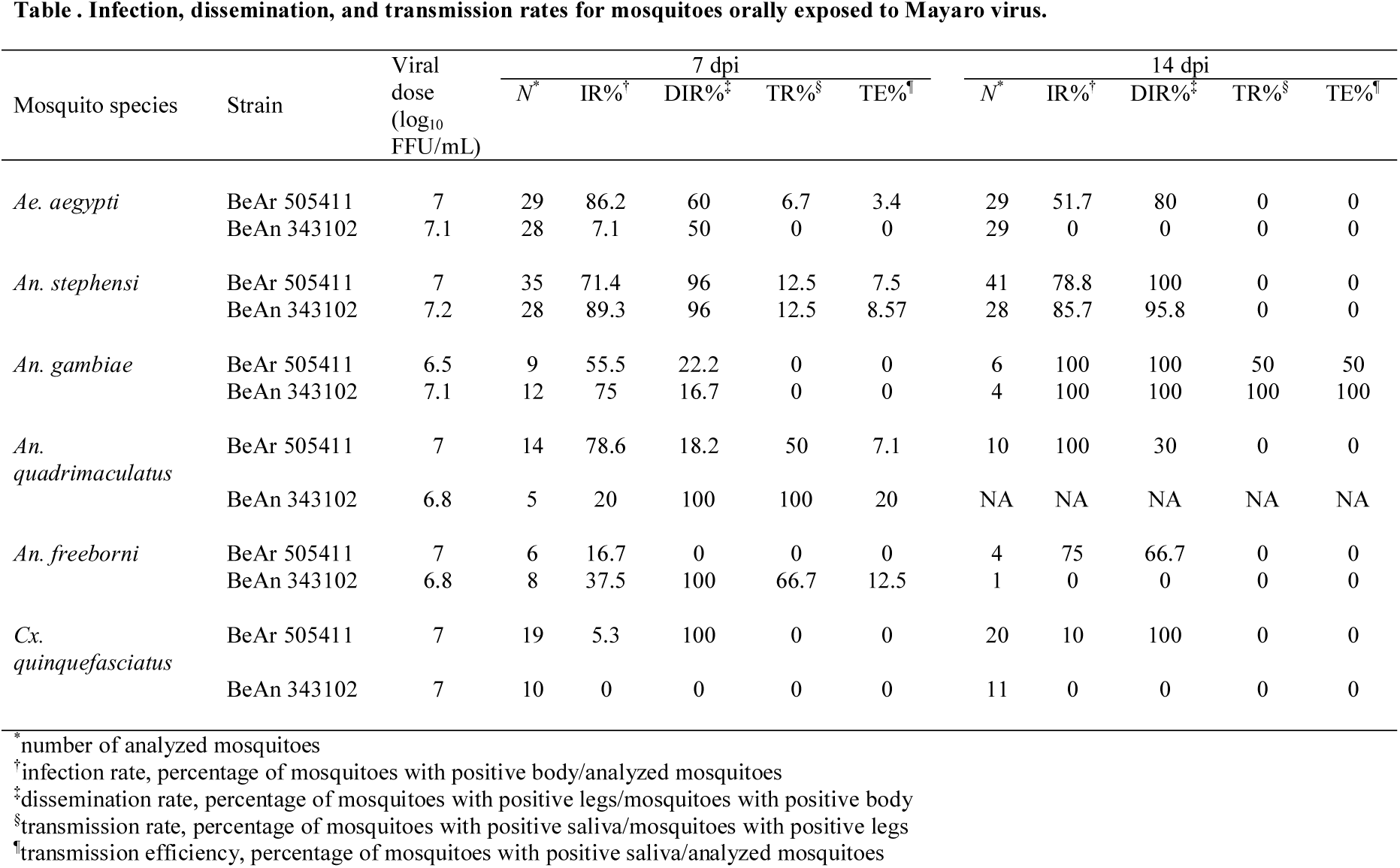
Infection, dissemination, and transmission rates for mosquitoes orally exposed to Mayaro virus.

At 7 and 14 dpi, mosquitoes were anesthetized with triethylamine (Sigma, St. Louis, MO, USA). Legs were detached from each body and placed in 2-mL tubes filled with 1 mL of mosquito diluent (20% heat-inactivated fetal bovine serum [FBS] in Dulbecco’s phosphate-buffered saline [PBS], 50 µg/mL penicillin/streptomycin, 50 µg/mL gentamicin, and 2.5 µg/mL fungizone) and a single zinc-plated, steel, 4.5-mm bead (Daisy, Rogers, AR, USA), and tubes immediately placed on ice. Saliva was collected using a capillary technique as previously described (*20*), expelled into in a 2-mL tube filled with 100 µL of mosquito diluent, and immediately placed on ice. Body and leg samples were homogenized at 30□ Hz for 2□ mi using a TissueLyser II (QIAGEN GmbH, Hilden, Germany) and centrifuged for 30 sec at 11,000 rpm. All samples were stored at −70°C until tested.

The presence of infectious MAYV particles in the body, legs, and saliva samples was tested by FFU assay in Vero cells. Vero cells were grown to a confluent monolayer in 96-well plates at 37°C with 5% CO_2_ in complete media (1× Dulbecco’s modified-essential media [DMEM], 100 units/mL penicillin/streptomycin, and 10% FBS). The next day, wells were washed with DMEM without FBS and incubated with a 30-µL aliquot of each homogenized tissue sample for 2 h at 37°C. After the incubation step, the 30-µL aliquot was removed from the cell monolayer and any unattached viral particles were removed with a DMEM wash. A total of 100 µL of overlay medium (1% methyl cellulose in complete growth medium) was dispensed into each well, and plates were incubated at 37°C inside the CO_2_ incubator. Cells were fixed at 24 h (bodies and legs samples) or 48 h (saliva samples) post-infection with 4% paraformaldehyde (Sigma, St. Louis, MO, USA). Fixed cells were blocked and permeabilized for 30 min with blocking solution containing detergent (3% bovine serum albumin and 0.05% Tween 20 in PBS) and washed with cold PBS. Viral antigens in infected cells were labeled using the monoclonal anti-Chikungunya virus E2 envelope glycoprotein clone CHK-48 (which reacts with Alphaviruses) (BEI Resources, Manassas, VA, USA) diluted 1:500 in blocking solution. Subsequently, cells were washed 4 times with cold PBS to remove unbound primary antibodies. The primary antibody was labeled with the Alexa-488 goat anti-mouse IgG secondary antibody (Invitrogen, Life Science, Eugene OR, USA) at a dilution of 1:500, and green fluorescence was observed and evaluated with an Olympus BX41 inverted microscope equipped with an UPlanFI 4× objective and a FITC filter (Figure 1). Fluorescent foci were counted for each well, and virus titers were calculated and expressed as FFU/mL.

**Figure 1.**
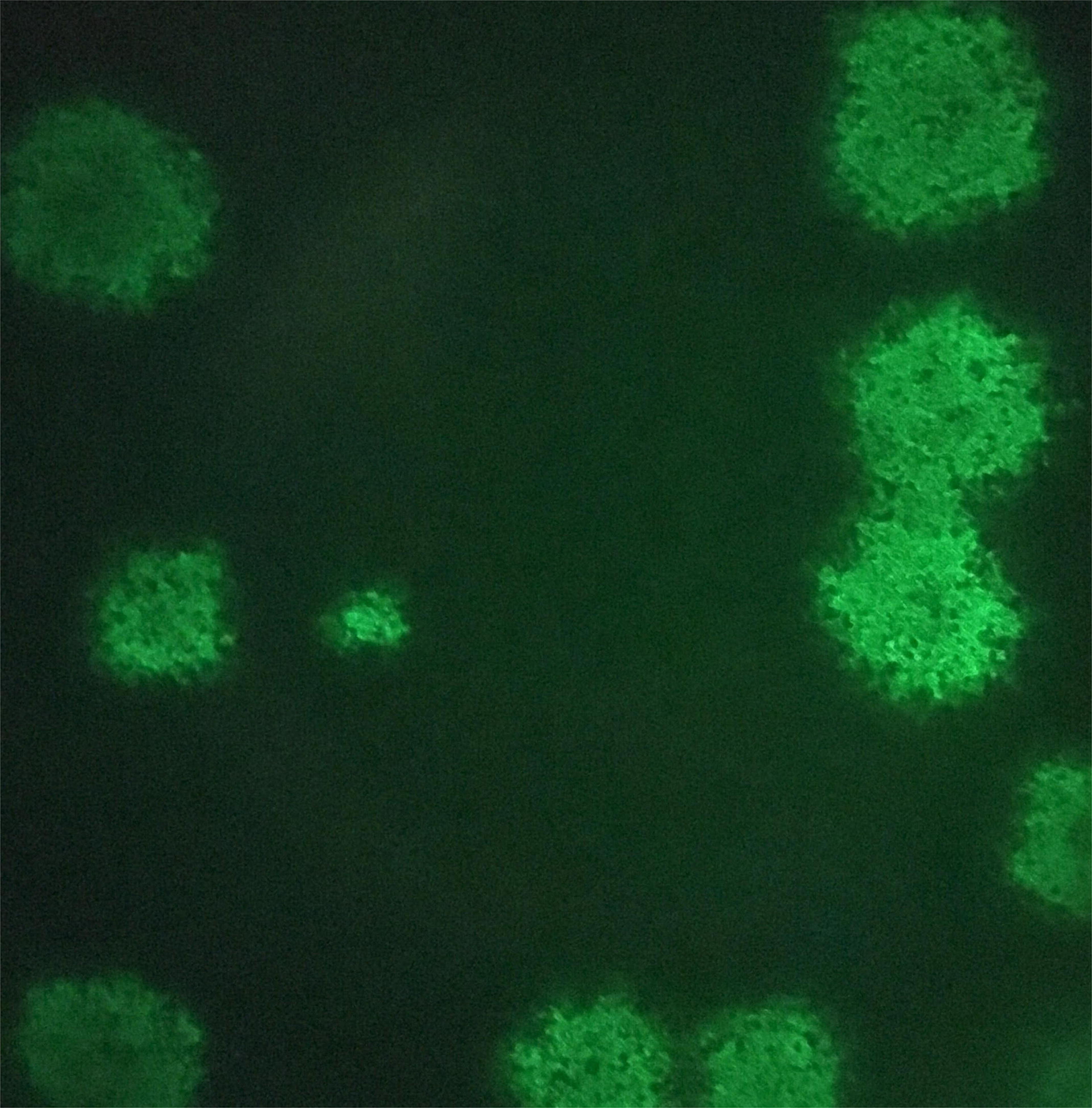
Fluorescent foci formation in Vero cells following infection with Mayaro virus. Vero cells were incubated with samples homogenate (body, legs or saliva). After 24 (body and legs) or 48 hours post infection (saliva) the monolayer was fixed, permeabilized, stained with antibody for alphavirus-E2 and visualized by immunofluorescence microscopy.

Data were analyzed using GraphPad Prism version 7.04. Differences in the IR, DIR, TR, and TE of mosquitoes challenged with BeAr 505411 and BeAn 343102 were analyzed by Fisher’s exact test. The Mann-Whitney U test was used to compare the body, legs, and saliva viral titers of mosquitoes exposed to BeAr 505411 or BeAn 343102.

## Results

A total of 115 *Ae. aegypti*, 132 *An. stephensi*, 31 *An. gambiae*, 29 *An. quadrimaculatus*, 19 *An. freeborni*, and 60 *Cx. quinquefasciatus* were analyzed in this study. Details of analyzed mosquitoes and the IR, DIR, TR, and TE are in Table 1.

All six mosquito species were susceptible to infection with MAYV to some degree, although there were MAYV strain–specific differences. IRs for *Ae. aegypti* exposed to strain BeAr 505411 were significantly higher compared to strain BeAn 343102 (p<0.0001) at 7 dpi, and IRs for strain BeAr 505411 at 7 dpi were significantly higher than 14 dpi (p<0.0001). Moreover, no *Ae. aegypti* exposed to strain BeAn 343102 became infected at 14 dpi despite the presence of positive mosquitoes at 7 dpi. IRs for *An. stephensi* and *An. gambiae* were similar across MAYV strains, and IRs increased over time in *An. gambiae*. *An. quadrimaculatus* and *An. freeborni* were susceptible to infection with both strains of MAYV, and *Cx. quinquefasciatus* was susceptible only to a low-frequency infection with strain BeAr 505411.

Once infected, all tested mosquito species developed a disseminated infection. Disseminated infection was generally detected as early as 7 dpi, with the exception of *An. freeborni* exposed to the BeAr 505411 strain. DIRs were similar for both virus strains in *An. stephensi* and *An. Gambiae* at both timepoints and for *Ae. aegypti* at 7 dpi. There was a trend toward higher DIRs for strain BeAn 343102 compared to strain BeAr 505411 in *An. quadrimaculatus* and *An. freeborni* at day 7. There was also a trend toward a higher DIR at 14 dpi than at 7 dpi for strain BeAr 505411 in *Ae. aegypti*, both strains in *An. gambiae*, and strain BeAr 505411 in *An. freeborni*.

Transmission was detected in all *Anopheles* species and *Ae. aegypti* (albeit very poorly), but not in *Cx. quinquefasciatus*. *An. stephensi*, *An. gambiae*, and *An. quadrimaculatus* were able to transmit both MAYV strains tested. For *Ae. aegypti* only a single transmission event was detected for virus strain BeAr 505411. Only virus strain BeAn 343102 was transmitted by *An. freeborni*. Both virus strains were able to be transmitted by *An. gambiae, stephensi,* and *quadrimaculatus*.

MAYV titers for all samples were calculated and expressed as FFU/mL (Figure 2). *Ae. aegypti* exposed to strain BeAr 505411 had significantly greater titers in the bodies (7 and 14 dpi) and legs (7 and 14 dpi) compared to strain BeAn 343102 (p<0.0001) (Figure). Conversely, *An. stephensi* exposed to strain BeAn 343102 had significantly greater titers in the bodies (7 dpi, p<0.05; 14 dpi, p<0.001) and legs (7 dpi, p<0.001) compared to strain BeAr 505411 (Figure 2). There were no significant differences in body, legs, or saliva titers between the MAYV strains in *An. gambiae*, *An. quadrimaculatus*, *An. freeborni*, and *Cx. quinquefasciatus*.

**Figure 2.**
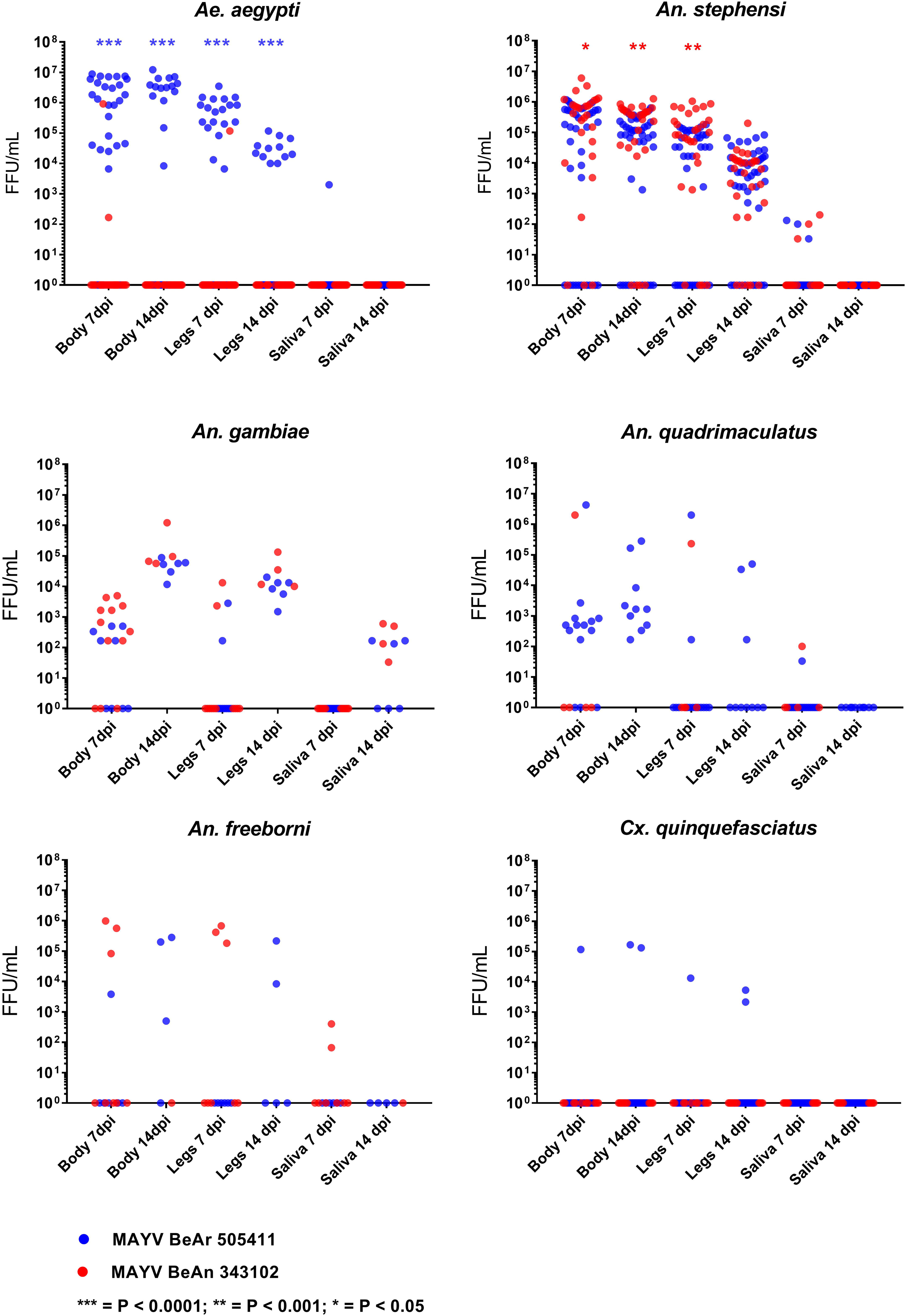
Viral titer in body, legs, and saliva of six mosquito species mosquitoes exposed to Mayaro virus. Each dot corresponds to a single mosquito sample. Viral titers were statistically compared between strains by Mann-Whitney U test.

## Discussion

These results demonstrate that *An. stephensi*, *An. gambiae*, *An. quadrimaculatus*, and *An. freeborni* are competent laboratory vectors for MAYV. The two viral strains tested present significant differences in their ability to infect and disseminate in *Ae. aegypti* and *An. stephensi*. In *An. stephensi*, the strain BeAn 343102 had a statistically higher titer in body and legs samples than BeAr 505411. Conversely, strain BeAn 343102 has a statistically lower body titer in *Ae. aegypti* and was not detected in legs, likely indicating the presence of a midgut escape barrier. Finally, strain BeAn 343102 failed to infect *Cx. quinquefasciatus*, likely due to the presence of a midgut infection barrier. The VC differences between the strains may be explained by the theory of the host genotype and pathogen genotype (G × G) interaction (*21*). G × G interactions have been found in many systems, including DENV (*22*). For example, Lambrechts et al. (*22*) showed that DENV vector competence varied greatly depending on the specific *Ae. aegypti* population and DENV genotype combination. This provides evidence that G × G interactions may be responsible for the adaptation of a lineage/strain to a specific population.

*Ae. aegypti*, *An. stephensi*, *An. quadrimaculatus*, and *An. freeborni* were able to transmit the virus at 7 dpi but we did not detect transmission at 14 dpi. The short extrinsic incubation period (EIP) of MAYV for these species might represent a notable increase in their vectorial capacity (*23*) and must be considered when establishing a future surveillance plan. In *An. stephensi*, the absence of transmission at 14 dpi corresponds with a decrease of the viral titer in the legs between 7 dpi and 14 dpi. These data suggest that in *An. stephensi*, MAYV infection may not persist, and may be progressively eliminated or limited by the vector. Similar results were recently published for *Ae. aegypti* infected with DENV (*24*). In that study, a progressive decrease of transmission began at 14 dpi and continued until 25 dpi, at which point no viral transmission was recorded. To test this hypothesis and to better understand the kinetics of MAYV infection, a study with a longer EIP and more intermediate timepoints should be performed. *An. gambiae* also is a competent laboratory vector for MAYV but the longer EIP (14 dpi) required for the transmission of the virus might limit the role of this species in the transmission cycle.

With the *Ae. aegypti* strain tested here, we obtained similar IR and DIR results compared to those previously described with a different strain (*16*). However, the MAYV TR in this study is considerably lower than that described by Long et al. (*14*) (6.7% vs. 88%). This discrepancy could be due to the genetic differences in the mosquito population (salivary gland infection barrier in the strain tested) or in the viral strain used for the experiment.

The global expansion of CHIKV due to a single point mutation (*30*) has previously demonstrated that the adaptation of an arbovirus to a new vector species can be devastating. The adaptation of MAYV to the *Aedes* vector has been analyzed (*31*), and the emergence of hybrid genotypes D and L suggests that *Aedes* mosquitoes can play an important role in the urban diffusion of MAYV. *Ae. aegypti* and *Ae. albopictus* are well adapted to urban and peri-urban habitats, and, contrary to *Haemagogus* mosquitoes, consistently have anthropophilic feeding behavior. Our results confirm that *Ae. aegypti* is a possible (if potentially inefficient) vector for MAYV, but more studies are needed to understand the differences in the VC for the genotype D and genotype L strains.

We found that *Cx. quinquefasciatus* can be infected with MAYV strain BeAr 505411, but is not able to transmit the virus. Conversely, another study found MAYV-positive *Cx. quinquefasciatus* during an outbreak of DENV in Mato Grosso, Brazil, and suggested that this species could sustain the transmission cycle (*15*). These results highlight the important point that merely detecting virus in a mosquito does not necessarily implicate it as a vector.

Previously, only two alphaviruses were known to be transmitted by Anopheles mosquitoes: O’nyong-nyong virus (*25*) and a single record for CHIKV (*26*). However, in the original paper describing the isolation of MAYV, the authors present an anecdote (no data) stating that when inoculated into *An. quadrimaculatus* from Trinidad, MAYV was able to replicate (although neither oral infection nor transmission was investigated) *(1)*. The capacity of *An. quadrimaculatus* and *An. freeborni* to transmit MAYV is particularly relevant to the United States, because the estimated geographic distribution of these species covers the entirety of the country (*27,28*). If MAYV was introduced into the United States, these two mosquito species may have the capacity to sustain the transmission cycle and spread the virus throughout the country. An interesting and important aspect of *Anopheles* vector biology is their tendency to have multiple feeding events during a single gonotrophic cycle (*29*). The bite frequency of *Anopheles* mosquitos increases their vectorial capacity and make them a very effective vector (*23*). For these reasons, we highlight the need for more studies on the possible role of *Anopheles* mosquitoes in spreading arboviruses in the United States.

We tested 4 Anopheles species (2 from North America, one from Africa, and one from Southeast Asia) for MAYV VC, and all were able to transmit the virus. Our results illustrate the knowledge gaps that remain about this important emerging virus. *Anopheles* mosquitoes in general are currently neglected as potential vectors of arboviral pathogens. Our data suggest that *Anopheles* sp. may be important vectors driving the emergence and invasion of MAYV (and potentially other arboviruses) across geographically diverse regions of the globe, and their epidemiological role in virus invasions should be further studied.

## Acknowledgments

We thank Ms. Erona Ibroci for assistance with mosquito rearing. This research was supported by NIH grants R01AI116636, R01AI128201, R21AI128918, and U19AI089672.

